# Horizontal Gene Transfer of Functional Type VI Killing Genes by Natural Transformation

**DOI:** 10.1101/122499

**Authors:** Jacob Thomas, Samit S. Watve, William C. Ratcliff, Brian K. Hammer

## Abstract

Horizontal gene transfer can have profound effects on bacterial evolution by allowing individuals to rapidly acquire adaptive traits that shape their strategies for competition. One strategy for intermicrobial antagonism often used by Proteobacteria is the genetically-encoded contact-dependent Type VI secretion system (T6SS); a weapon used to kill heteroclonal neighbors by direct injection of toxic effectors. Here, we experimentally demonstrate that *Vibrio cholerae* can acquire new T6SS effector genes via horizontal transfer and utilize them to kill neighboring cells. Replacement of one or more parental alleles with novel effectors allows the recombinant strain to dramatically outcompete its parent. Through spatially-explicit simulation modeling, we show that the HGT is risky: transformation brings a cell into conflict with its former clonemates, but can be adaptive when superior T6SS alleles are acquired. More generally, we find that these costs and benefits are not symmetric, and that high rates of HGT can act as hedge against competitors with unpredictable T6SS efficacy. We conclude that antagonism and horizontal transfer drive successive rounds of weapons-optimization and selective sweeps, dynamically shaping the composition of microbial communities.

## Introduction

Horizontal gene transfer (HGT) by plasmid conjugation, viral transduction, and by natural transformation plays a fundamental role in the evolution of bacteria, archaea, and also in plants and other eukaryotes [1]. Genomic analyses implicate HGT as a major factor responsible for the mosaic genomes of many bacteria, including the waterborne microbe *Vibrio cholerae*, which often carries the cholera toxin (CTX)-encoding prophage responsible for major cholera outbreaks in Haiti and endemic regions [2-5]. *V. cholerae* isolates from aquatic environments are invariably non-toxigenic (CTX^-^) and commonly associated with chitinous surfaces like shells of crabs or zooplankton that may promote transmission [6].

All sequenced isolates of *V. cholerae* encode a type VI secretion system (T6SS) weapon that can deliver toxic effector proteins directly into grazing predators, diverse proteobacteria, and other *V. cholerae* that lack cognate immunity proteins (reviewed in [7-11]). The majority of non-clinical isolates express the T6SS constitutively, in the absence of chitin, likely as a competitive strategy in complex microbial communities of environmental biofilms [12, 13]. By contrast, the majority of toxigenic clinical *V. cholerae* isolates engage in contact-dependent T6-killing only when triggered by the presence of chitin [14, 15], perhaps down-regulating this activity in a host [9]. Interestingly, chitin can also induce natural transformation in clinical isolates, which can promote HGT events due to lysis of adjacent non-immune neighbors [14]. Genome analyses of *V. cholerae* and other *Vibrio* species suggest that T6 loci encoding distinct effectors and adjacent cognate immunity factors, yet flanked by highly conserved sequence, may themselves be horizontally acquired by homologous recombination [16-18]. Furthermore, it was hypothesized, but not experimentally tested, that HGT of distinct effector/immunity pairs could enable populations to diversify by generating individuals compatible with their competitor, yet susceptible to kin [16, 18]. Whether HGT of novel T6 genes is adaptive in communities where kin and non-kin are in close proximity is unclear. Most clinical *V. cholerae* strains, which constitute the majority of sequenced and characterized isolates, carry identical T6 loci, thus hampering efforts to address this challenge.

We recently demonstrated that two sequenced *V. cholerae* strains with two syntenic T6 loci encoding distinct effector/immunity pairs behaved as “mutual killers” that precipitate a phase separation when co-cultured [19]. One strain is derived from clinical reference isolate C6706 that is induced by chitin to express both the DNA uptake apparatus and the T6SS. The other is recently sequenced environmental isolate 692-79 [20], defective at DNA uptake and yet T6SS-proficient. Here we show that co-culturing of these two strains in the presence of chitin induces unidirectional horizontal transfer of novel T6 effector/immunity pairs to the clinical isolate by homologous recombination. Transformants are simultaneously protected from attack by the environmental donor and susceptible to attack by former siblings. Modeling confirms that the fitness costs and benefits from acquisition and replacement of distinct T6SS effectors are contextual and depend on spatial structure of the population, relative efficacy of T6SS-effectors, and the rate of horizontal transfer. These results reveal that members of microbial communities can continuously adapt to diverse competitors by rapid acquisition of new molecular weaponry via horizontal transfer.

## Results

The core T6SS components of all sequenced *V. cholerae* strains are encoded in three operons – one main cluster and two auxiliary clusters Aux1 and Aux2 (Fig. 1), while additional T6SS loci are occasionally present that may encode new effector/immunity pairs thought to be dispensable for assembly of the T6SS spike [18, 21]. Each operon terminates in a gene encoding a toxic effector, immediately followed by its cognate immunity gene. In the toxigenic C6706 strain, the main cluster effector protein, VgrG-3, is a ‘specialized’ effector that forms a physical component of the puncturing spike tip, while the Aux1 and Aux2 effectors are ‘cargo’ effectors that each require a specific Tap/PAAR chaperone for assembly with its cognate VgrG spike protein [8, 22]. Aux1 and 2 operons of all sequenced *V. cholerae* strains have conserved DNA sequences flanking a variable region of differing %GC content that encodes the effector-immunity pair, Tap, and C-terminus of VgrG [18]. The main T6SS cluster is entirely conserved between *V. cholerae* strains except for a short VgrG-3 C-terminus and a portion of the immunity gene.

**Fig. 1.**
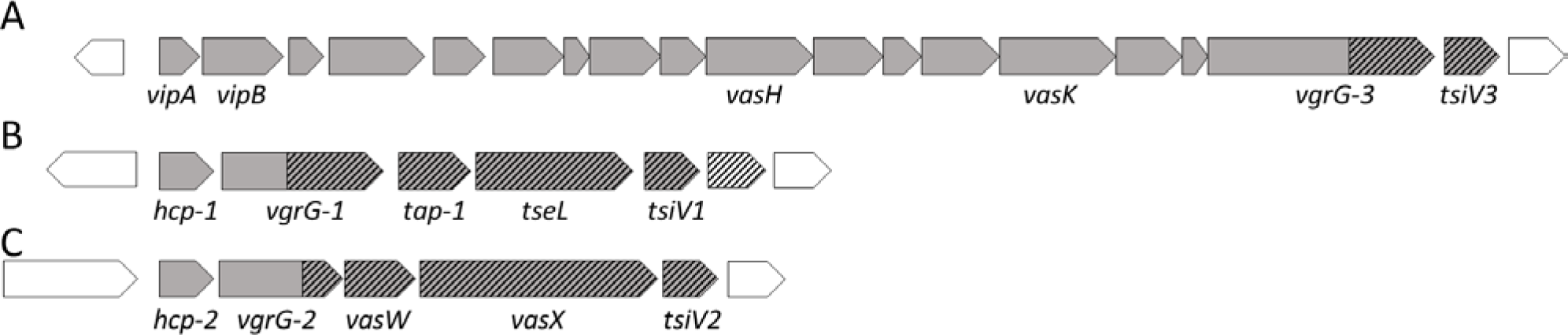
Organization of T6SS operons in *Vibrio cholerae* C6706. T6SS genes, shaded in grey, are organized into one large cluster (A) and two auxiliary clusters Aux1 (B) and Aux2 (C). Hatched regions that are variable when compared to other sequenced *V. cholerae* isolates. The first gene of each of the auxiliary clusters encodes the major subunit of the protein tube, Hcp. Genes encoding effector and immunity proteins are respectively, *vgrG-3* and *tsiV3* (main cluster), *tseL* and *tsiV1* (Aux-1) and *vasX* and *tsiV2* (Aux-2).

As genomic analyses predict that transfer of entire T6SS operons frequently occurs in nature [16, 18], we tested this experimentally with toxigenic reference strain C6706, which is induced by chitin to become naturally transformable, and a nontransformable environmental isolate 692-79 [12]. At the Aux1 locus, both strains have distinct phospholipase effectors; TseL in C6706 and a phospholipase similar to Tle1 of *P. aeruginosa* in 692-79 [23, 24]. The Aux2 operons of the two strains encode effectors with entirely different activities: a VasX pore forming protein in C6706 and a LysM domain-containing effector that potentially targets peptidoglycan in 692-79 [25, 26]. We recently demonstrated that the distinct T6 activities of these strains allowed them to engage in mutually antagonistic T6S killing [19]. We therefore predicted that horizontal transfer of T6SS genes via natural transformation from a 692-79 donor to a C6706 recipient would generate transformants with T6SS profiles distinct from both donor and recipient strains. To test whether we could observe C6706 transformants that acquired the Aux alleles of the 692-79 donor, we introduced antibiotic resistance cassettes by recombination directly downstream of the Aux1 or Aux2 immunity genes in the 692-79 strain. To avoid challenges interpreting exchange of core and regulatory components, we did not design a system to measure exchange of large cluster components. When the two strains were co-cultured on chitin tiles submerged in artificial seawater, C6706 acquired the entire Aux1 or Aux2 operons of 692-79 by natural transformation at frequencies of ∼1×10^−6^, while a nontransformable C6706 mutant lacking the *comEA* gene did not (Fig. S1). Acquisition of each T6SS operon was confirmed by PCR analysis. C6706 cultured with chitin was able to acquire both Aux1 and Aux2 T6SS alleles either from genomic DNA derived from 692-79 or by co-culture with the donor strain (Fig. S1) and successive rounds of chitin-induced transformation also generated a C6706 transformant with both Aux1 and Aux2 allele replacements. Since acquisition by homologous recombination of a new T6SS operon is accompanied by a simultaneous loss of the original alleles, we initially expected that transformants would be selected against due to T6S-killing by neighboring former clonemates. However, T6SS-inactivation by deletion of *vasK* encoding an assembly protein [27] had modest effects on acquisition frequency of new T6SS operons either when using genomic donor DNA or in co-culture (Fig. S1). We suspect this is likely because the substrate adhered cell-cell contact required for T6SS-mediated killing is not consistently maintained in the seawater conditions used in our transformation assays.

Recombination of each entire Aux cluster in the C6706 transformants was verified by sequencing the effector and immunity genes and a stretch of DNA 2 kilobases upstream of the start codon of the first gene in each operon. Sequencing confirmed acquisition of both the effector and immunity genes of the 692-79 donor, but newly acquired T6SS effectors must be expressed, properly folded, and loaded onto the T6SS complex encoded by the recipient strain if they are to be functional. Indeed we found that the C6706 transformants with the Aux1 T6SS operon from 692-79 (which we refer to here as A1), the Aux2 operon (A2), and both operons (A1,2) are able to kill C6706 to an increasing degree in standard 3-hour T6 killing assays (Fig. 2A). Acquisition of either cluster also diminished the ability of transformants to kill 692-79, however, replacement of both clusters did not completely abrogate killing of 692-79 (Fig. 2B). We predicted that the residual killing activity might be due to expression of the main cluster VgrG-3 of C6706, which was not replaced in these experiments. Indeed, deletion of *vgrG3* in the double (A1,2) transformant completely abolished killing of the donor 692-79 (Fig. S2). Replacement of T6SS clusters also changed the immunity profiles of each strain: a single or double Aux cluster replacement protected the transformant from killing by the 692-79 donor (Fig. 2C) but made the strains susceptible to killing by the C6706 recipient (parent), with the largest effect seen in the Aux1,2 transformant (Fig. 2D).

**Fig. 2.**
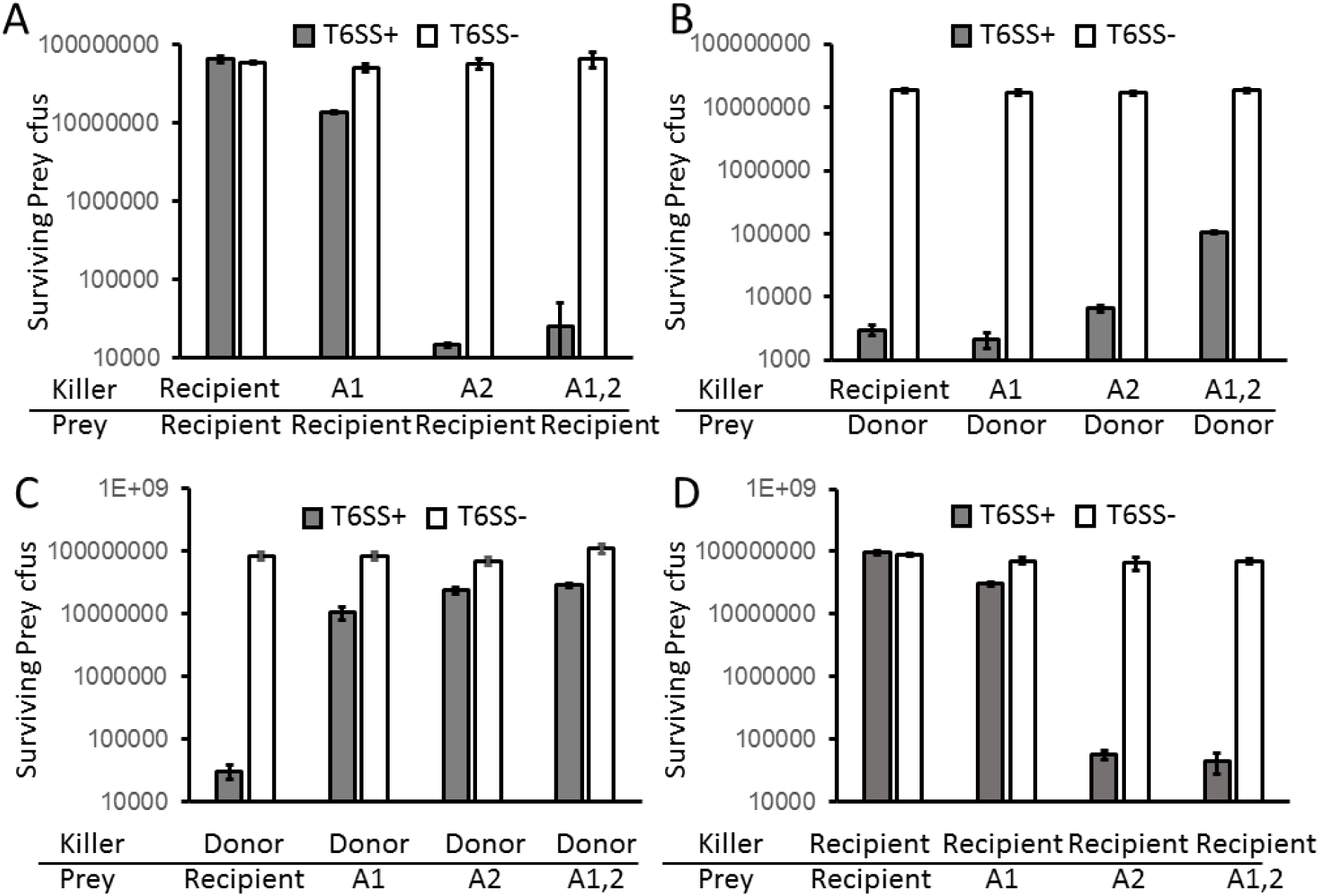
Horizontally acquired T6SS effectors and immunity proteins are functional in the recipient strain. ‘Killer’ strains and T6SS^−^ Δ*vasK* ‘prey’ strains were mixed at a 10:1 ratio, incubated at 37C for three hours on membrane filters and surviving prey CFUs were counted on selective media. Killer strains were either T6SS^+^ (grey bars) or T6SS^−^ (Δ*vasK*, white bars). Strains used were C6706 (recipient), 692-79 (donor) and C6706 derivatives in which either Aux1 (A1), Aux2 (A2) or both clusters (A1,2) were replaced by those of 692-79.

A previous study demonstrated that the effector-immunity alleles carried by *Vibrio cholerae* V52 (and C6706) were the most potent in pairwise competition experiments amongst the different allele combinations tested [18].To examine the relative fitness of each T6SS transformant with respect to the recipient (parent) and donor, we performed pairwise 24h competition experiments (Fig. 3). Acquisition of each T6SS cluster conferred a significant advantage to the transformant relative to the recipient (Fig. 3A). Each transformant carrying an Aux1 (A1), Aux2 (A2) or both Aux1 and 2 (A1,2) replacement has increased fitness compared to both recipient and donor and this fitness gain is T6SS-dependent (Fig. 3A-C). The A1,2 transformant has a competitive advantage over every other strain tested, suggesting that when 692-79 is the donor, acquisition of T6SS alleles by HGT is always beneficial to the C6706 recipient. However, the donor has a large T6SS-independent fitness defect compared to C6706 and its derivatives (Fig. 3C) and represents a potentially rare example of a poor competitor that carries superior weapons. To verify the competitive hierarchy we observed in two-partner competitions (Fig. 3D), we co-cultured C6706 and each transformant (A1, A2 and A1,2) in an initial 1:1:1:1 ratio for 5 days with back dilution and repeat passaging every 24 hours (Fig. S3). We found that the A1,2 transformant dominated the mixed culture within 24 hours and the relative representation of the other strains was in agreement with the hierarchy observed in two partner competitions (Fig. 3). Thus, HGT of T6SS allows C6706 to rapidly improve its repertoire of T6SS weaponry.

**Fig. 3.**
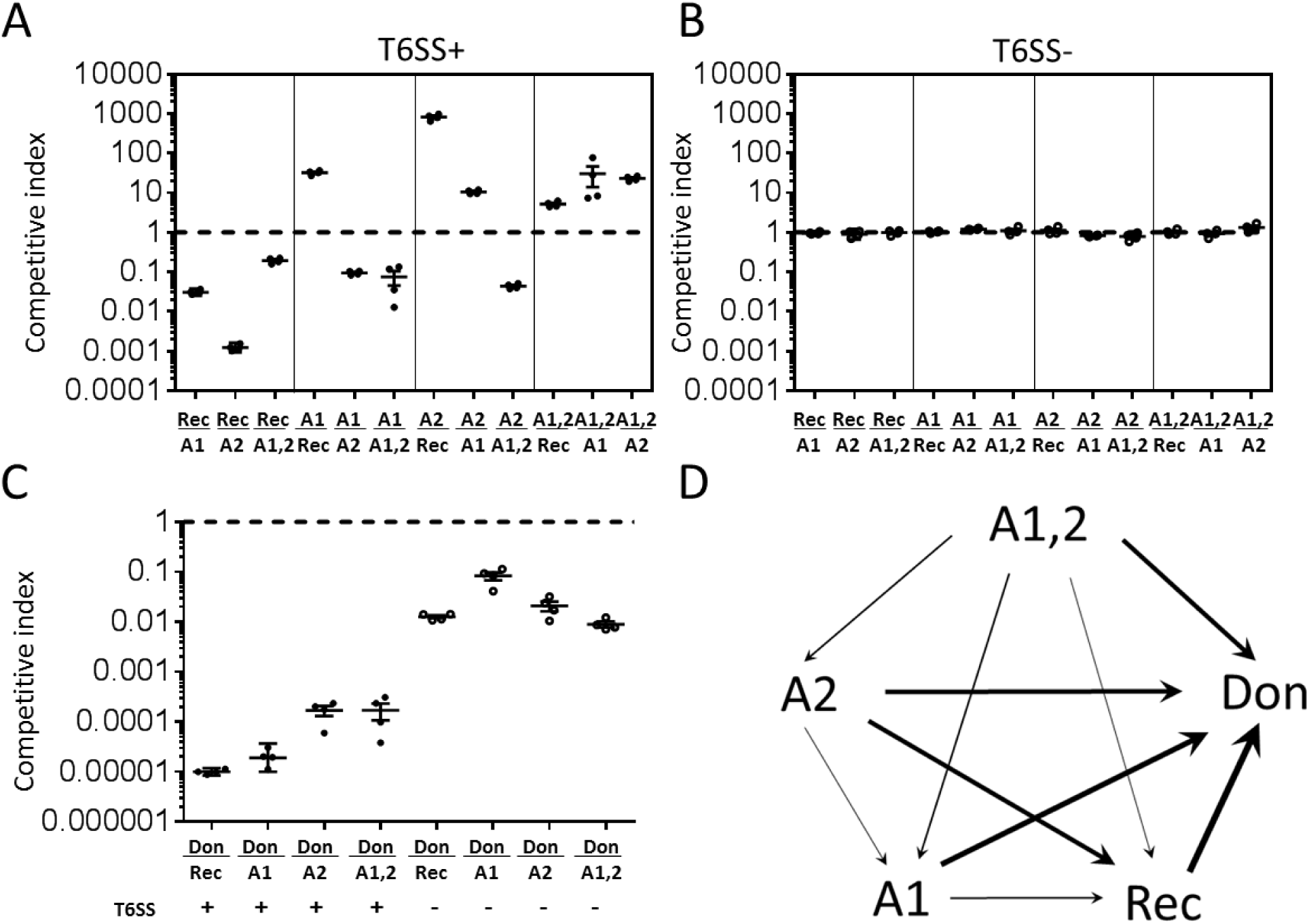
Acquisition of T6SS effectors from a donor (692-79) is beneficial to the recipient (C6706). (A and B) Pairwise competitions were set up between the recipient strain C6706 (Rec), the donor 692-79 (Don) and C6706 derivatives in which either Aux-1 (A1), Aux-2 (A2) or both clusters (A1,2) were replaced by those of 692-79. Strains were mixed at a 1:1 ratio and incubated on solid agar in a 12-well plate at 37C for 24 hours and survivors of each strain were counted on selective media. Both strains were T6SS+ (panel A) or T6SS− (Δ*vasK*, panel B). Competitive indices are measured as the final ratio of survivors divided by the initial ratio of inoculation. Horizontal lines in each cluster indicate geometric mean of competitive indices. (C) Competitive hierarchy of strains analyzed in A and B is represented with arrows proceeding from the more fit competitor towards the less fit competitor. Line weight of each arrow is proportional to the log_10_ of the geometric mean of the competitive indices of the [more fit/less fit] competitor.

To derive general principles for how natural selection might act on the HGT of T6SS clusters, we extended a model we previously developed to consider growth and competition on a two dimensional lattice between any two mutually-antagonistic strains [19]. In our simulation, each strain possessed three distinct T6SS alleles. The ‘recipient’ strain was capable of sequentially replacing its T6 effectors with those obtained from the ‘competitor’ strain (see Methods for model details). As in our experiments, the recipient was capable of HGT, but the donor was not. We observed that HGT can have a profound impact on T6SS-mediated competition dynamics. When the recipient possesses inferior T6SS effectors, HGT events precipitate a series of selective sweeps resulting in transformants that have completely replaced their T6SS alleles with those obtained from the superior competitor strain (Supplementary Movie 1). As a result, HGT allows for competitive rescue of a strain that encodes an inferior set of T6SS alleles (relative T6SS efficiency <1). More generally, the consequences of HGT on the fitness of the recipient genotype depend on how its T6SS loci compare to its competitor’s, and the degree to which the population is spatially structured (Fig 4a relative to 4b). Higher HGT rates are favored when a competitor possesses superior T6SS alleles; but costly when the competitor’s T6SS alleles are inferior (Fig. 4). The benefits of HGT are far greater when the population is highly structured with each strain spatially separated (*e.g*., Figure 4a), since T6 cluster replacement via HGT takes time. When the population is highly structured, the superior competitor takes more time to displace the recipient, which allows sufficient time for HGT and allelic replacement of T6SS clusters. The fitness valley in Fig. 4a occurs when horizontally-acquired T6SS alleles are slightly better than those possessed by the recipient, allowing them to kill their clonemates, but not enough to escape their geometric disadvantage and spread. As a result, HGT drives high rates of intra-clonal conflict, reducing their relative fitness.

**Fig. 4.**
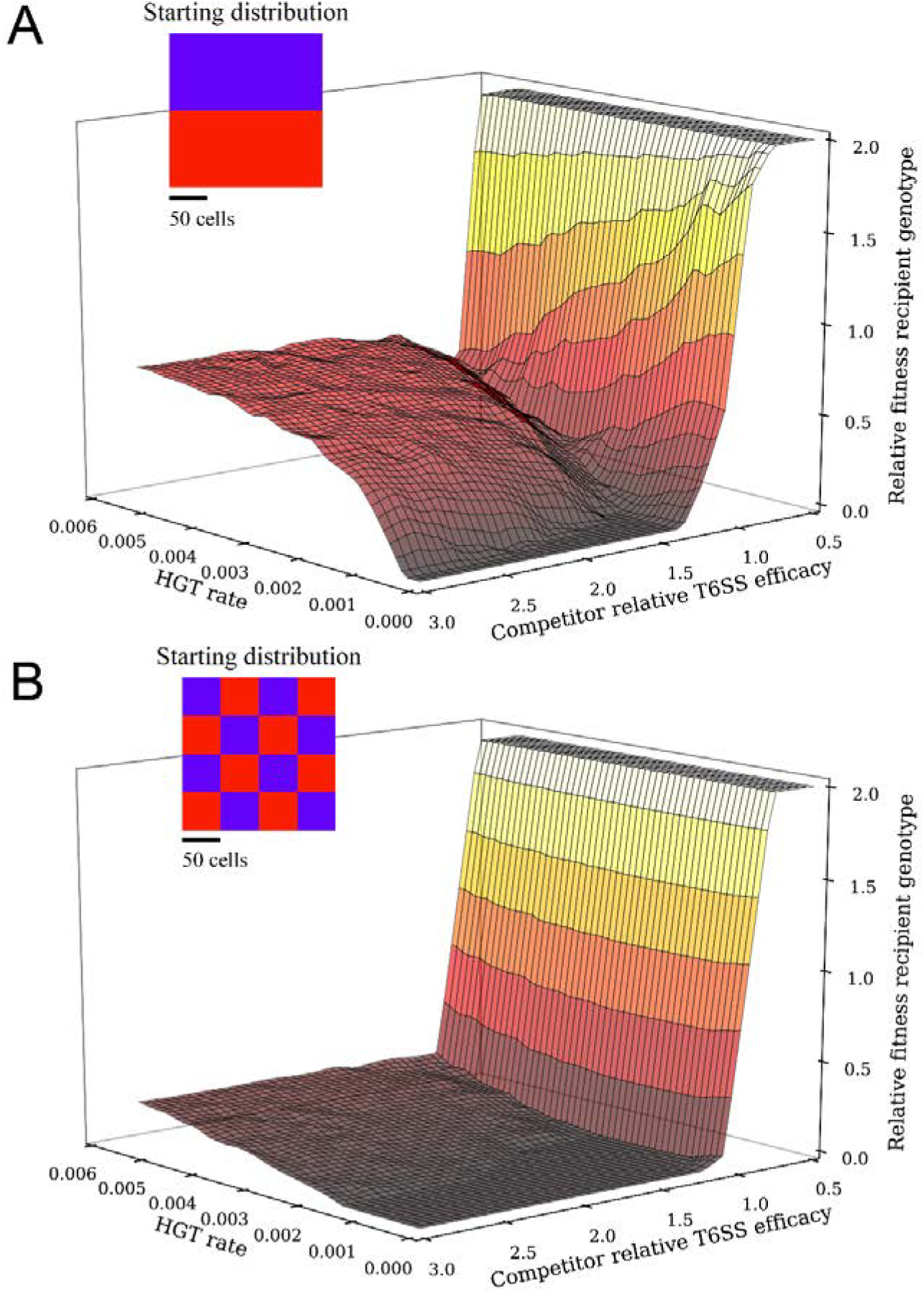
Costs and benefits of horizontally acquiring a competitor’s T6SS. In this model of competition between two bacterial strains, one (the ‘recipient’) was capable of integrating T6SS alleles from the other (the ‘competitor’). High rates of HGT were only adaptive when the recipient strain faced a competitor with superior T6SS alleles. For plotting purposes, relative fitness > 2 was clipped. Simulations were run for 15,000 time steps. Timelapses of these fitness landscapes are shown in Supplementary Movies 2 and 3.

T6SS-mediated conflict is inherently risky, as competitors vary widely in their relative killing efficacy [18]. While HGT may allow a strain with inferior T6SS effectors to survive an interaction with a superior competitor in which it might otherwise be eradicated, HGT may also be costly, as rare transformants that acquire foreign T6SS loci immediately turn on adjacent former clonemates and are usually killed (Supplementary Movie 1). While it is clear that incorporating *inferior* T6SS effectors will be costly, this is a separate cost from intra-strain genetic conflict, which occurs even when superior T6SS effectors are obtained horizontally. To estimate the fitness consequences of HGT-mediated intra-strain conflict, we simulated competition when mutually antagonistic recipients and competitors had distinct T6SS clusters that nonetheless killed with equal efficacy, and then varied HGT rate (Figure 5, blue points). We measured the geometric mean fitness, as this best predicts long-term changes in the frequency of each T6 effector, and is especially useful for taking variance in fitness across generations into account (Simons 2009). Within-strain conflict, which can be estimated by the regression of fitness on HGT rate when partners do not vary (blue points; y=−74.2x+0.991, r^2^=0.99), was very costly, reducing fitness by 40% when the per-cell probability of HGT per timepoint was 0.005.

**Fig 5.**
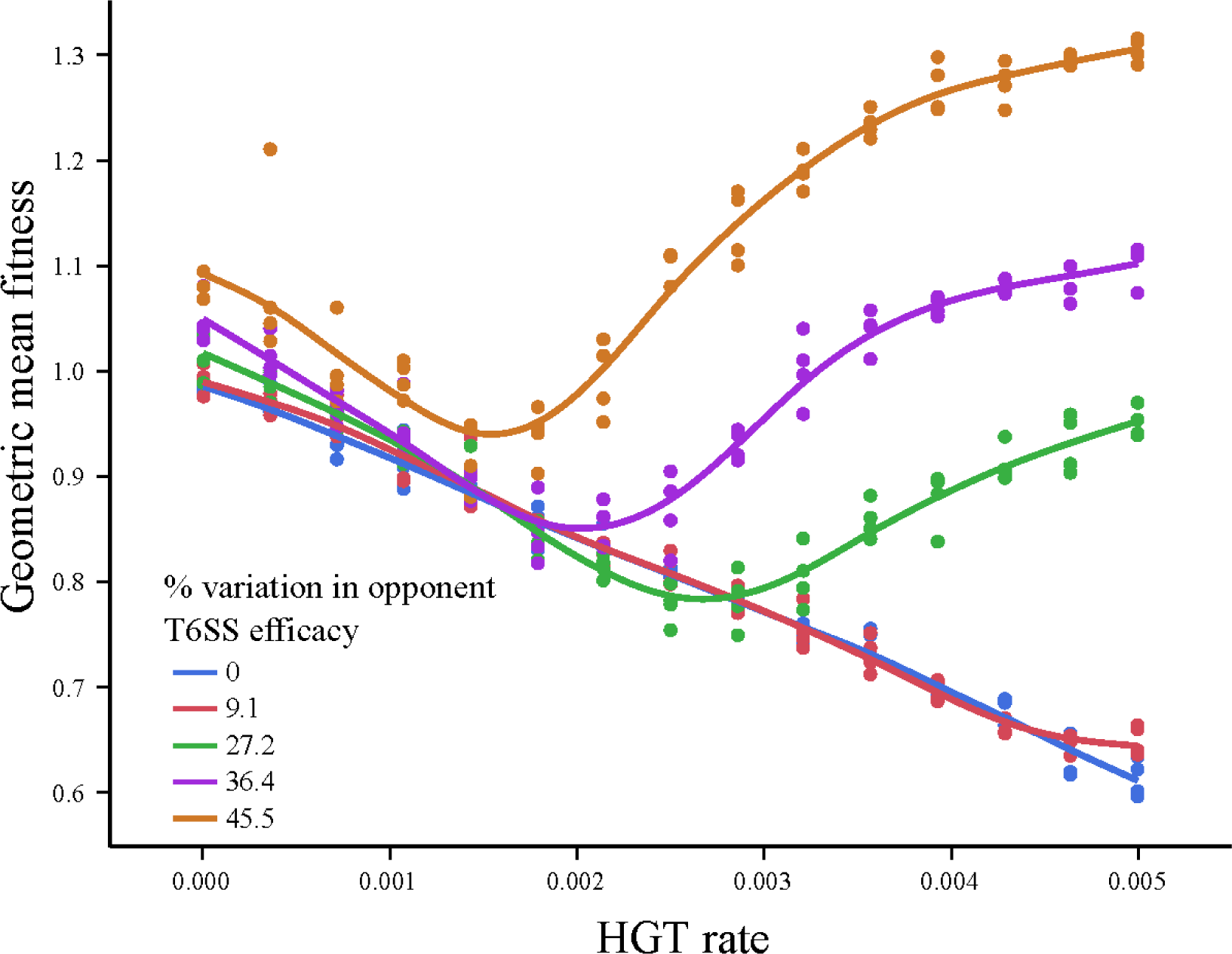
Horizontal gene transfer serves as a hedge against unpredictable competitors. To examine effects of variability in competitor T6SS efficacy on selection on HGT rate, the geometric mean fitness of the recipient genotype was determined across pairs of simulations, in which recipients faced competitors with increasingly high variation in T6SS efficacy (±0, 9.1, 27.2, 36.4, and 45.5% from its own T6SS).

The fact that HGT incurs significant costs due to intra-strain conflict and benefits via survival when facing a superior competitor raises the possibility that it may act as a hedge against uncertainty in the T6SS efficacy of future opponents. We examined this hypothesis by calculating a recipient strain’s geometric mean fitness across a pair of interactions in which it faced a competitor with symmetrically-varying T6SS efficacy (there was thus no overall bias in favor of either strain) and across a range (0-0.005) of HGT. Geometric mean fitness above 1 indicates that HGT is adaptive. When the two strains have T6SS with similar efficacy, the costs of HGT (intra-strain conflict) exceed its benefits and selection acts against HGT (Fig. 5). However, when competitor quality is highly variable, HGT can be adaptive, with selection favoring very high rates of transformation (Figure 5). The valley-shaped distribution of fitness for the high-variance treatments is a consequence of the geometry of competition: when HGT rates are low, transformants are surrounded by former clonemates that can now kill them via T6SS. Isolated transformants have a low probably of surviving killing attempts by their neighbors, even if they have moderately superior T6SS alleles. Higher rates of HGT lead to rare but important multiple transformation events where two or more adjacent cells uptake a competitor’s allele, reducing their geometric disadvantage and dramatically increasing the probability that they will precipitate a selective sweep. Note, even when HGT = 0, variation in competitor T6SS increases the recipient’s fitness (Figure 5). This reflects the cost of variability on the competitor’s fitness (or, inversely, the relative benefit the recipient experiences by not varying), a well-known phenomenon in evolutionary ecology [28]. Thus, high rates of HGT were strongly beneficial when the competitor’s T6SS efficacy was more variable, as the increased survival in the face of a superior competitor more than made up for HGT’s costs.

## Discussion

We have demonstrated that toxigenic bacteria can acquire T6SS effector-immunity pairs from non-toxigenic neighbors via natural transformation, consistent with the hypothesis that this frequently occurs in the environment [16, 18]. Once acquired, the novel T6SS effector-immunity proteins are functional and can contribute towards T6SS-mediated antagonism against neighboring cells as well as provide immunity against T6SS attacks by neighbors. Interestingly, transformants that undergo successive HGT of T6SS effector-immunity pairs have increased fitness relative to either parental strain in direct competition experiments, reaffirming the fundamental role of HGT in shaping bacterial ecology and evolution. However, our models show that the fitness consequences of T6SS acquisition are contextual, depending on the spatial structure of the population, rate of horizontal transfer, and the relative efficacy of the molecular weapons being acquired. Our modeling reveals that HGT can be quite costly: a cell runs the risk of incorporating an inferior T6SS allele, and every transformant must overcome the geometric disadvantage of being surrounded by former clonemates to which it is no longer fully immune. Still, these costs are more than compensated for when an HGT+ strain is facing a competitor with far superior T6SS alleles. Indeed, we show that variability in opponent T6SS allele quality alone can select for high rates of HGT, allowing HGT to act as a hedge against uncertain opponent quality.

In our experimental system, we were only able to observe examples of T6SS cluster replacement. However, bioinformatic and phylogenetic approaches have recently identified lineages of *V. cholerae* that appear to have undergone multiple rounds of HGT at a single highly recombinogenic T6SS locus to form ‘immunity gene arrays’ encoding several putative T6SS immunity proteins, which may provide protection against effectors from multiple *V. cholerae* and non*-V. cholerae* strains [16]. Successive acquisition of the immunity genes likely benefits the transforming lineage in three ways. First, a recipient strain gains immunity against a diverse set of T6SS+ competitors. Second, the recipient avoids the metabolic cost of expressing non-native effectors. Third, retention of the existing effector-immunity pair (as opposed to the replacement observed here) avoid costs of intra-strain genetic conflict. This represents an additional mechanism to promote horizontal spread of T6SS alleles while mitigating the costs of HGT.

In toxicogenic *V. cholerae* strains C6706 and A1552, both natural transformation and T6S are co-induced under conditions of chitin utilization, starvation, and quorum sensing at high cell density (Watve, Borgeaud). Specifically, activation of the quorum sensing system by *Vibrio*-specific autoinducer molecules induces the DNA uptake machinery [29, 30]. This concordance suggests that *V. cholerae* may have evolved to acquire genes, including T6S effector-immunity pairs, from recently killed competitors. This has, until now, presented a conundrum: why acquire genes from defeated, inferior competitors? Our work demonstrates that the acquisition of novel T6S systems provides an efficient route to means-test novel weaponry, which may be strongly favored by natural selection if competitor T6S efficacy varies widely through time. More generally, our results suggest that the interplay of antagonism and horizontal transfer results in successive rounds of combat, arms-trading, and selective sweeps that may dynamically shape diverse microbial communities.

## Materials and Methods

### Bacterial strains and culture conditions

All *V. cholerae* strains were derivatives of El Tor C6706 *str-2* [31] or environmental strain 692-79 (Table S1, supplemental material). Bacteria were routinely grown at 37°C in lysogeny broth (LB) under constant shaking, or statically on LB agar, supplemented with ampicillin (100 μg/mL), kanamycin (50 μg/mL), chloramphenicol (10 μg/mL for *V. cholerae* and 25 μg/mL for *E.coli*), spectinomycin (100 μg/mL), streptomycin (5 mg/mL), and diaminopimelic acid (50 μg/mL) where appropriate.

### Recombinant DNA techniques and construction of *V. cholerae* mutants

In-frame deletions and promoter-replacement mutants in *V. cholerae* were constructed by previously described allelic exchange methods [32]. Standard molecular biology-based methods were utilized for DNA manipulations. DNA modifying enzymes and restriction nucleases (Promega and New England Biolabs), Gibson assembly mix (New England Biolabs), Q5, Phusion and OneTaq DNA Polymerases (New England Biolabs) were used following the manufacturer’s instructions. All recombinant DNA constructs were tested by PCR and verified by Sanger sequencing (Eurofins).

### Transformation assays

Assays for natural transformation were performed as previously described [33]. Briefly, the recipient strain was back-diluted from an overnight liquid culture and grown at 37°C in LB broth with shaking until it reached an OD_600_ of 0.4-0.6. Cultures were washed in artificial seawater (ASW), diluted to OD_600_ of 0.15, and 2 mL cultures in ASW were incubated at 30°C with a chitin tile for 24 hours. The chitin tile with adherent bacteria was moved to fresh 2 mL ASW containing 1 μg/mL of genomic DNA marked with an antibiotic resistance cassette and incubated at 30°C for a further 24 hours. Cells were collected by washing the chitin tile in 2 mL of fresh ASW, and dilutions were plated on selective media to determine transformation frequency. Acquisition of T6SS alleles from donor DNA linked to the antibiotic resistance cassette was verified by allele specific multiplex PCR using primers specific for C6706 or 697-79. Transformation in co-culture was performed as described above with the following exceptions: both donor and recipient strain were each grown to an OD_600_ of 0.4-0.6 in LB broth, diluted to an OD_600_ of 0.15 after washing in ASW mixed in a 1:1 ratio and 2 mL of each mixed culture was inoculated with a chitin tile at 30°C. No exogenous genomic DNA was added.

### Killing assays

T6SS killing assays were performed as previously described [15]. Killer and prey strains grown overnight in LB broth at 37°C were centrifugated and resuspended in fresh medium to an OD_600_ of 1.00. Killer and prey strains were mixed at a ratio of 10:1 and 50 μL of each suspension was spotted onto a gridded filter disc (Whatman) placed on a LB agar plate and confined to 9 grid squares (3.1 × 3.1 mm each) to reduce variability. Filters were incubated at 37°C for three hours, washed with 5 mL LB broth to dislodge cells and dilutions were plated on selective media to determine prey survival.

### Pairwise competitions

Competitor strains were grown overnight in LB broth at 37°C were centrifugated and resuspended in fresh medium to an OD_600_ of 1.00. Pairs of strains were mixed 1:1 and 50 μL of each suspension was inoculated on LB agar plugs in a 12 well plate, dried under laminar airflow in a biosafety cabinet and incubated overnight at 37°C. Cells were resuspended in 1 mL LB broth and dilutions were plated on selective media to determine survival of each competitor. Competitive index was determined as [final ratio of survivors/initial ratio of inoculation].

### Individual-based simulation

We extended the individual-based model from McNally et al [19]. 200×200 cell square lattices were seeded with an equal number of ‘recipient’ and ‘competitor’ genotype cells. Biologically, this is analogous to a population growing across a solid surface, such as a crab shell. The starting conditions for Figure 4 are shown as insets; all other simulations use the well-separated population shown in Figure 4a. At each time point, 5% of the cells in the population were selected to reproduce if an adjacent space is unoccupied. Similarly, 5% of the cells in the population were selected to activate their T6SS. The probability that they kill a neighboring cell depended on the difference between their T6SS effector / immunity pairs, and the relative killing efficiency of those effectors. The three effectors possessed by each genotype were assumed to act in an additive manner, so the probability of killing was determined by summing across the number of effector / immunity pairs that differ between the strains, weighted by the efficacy of those effector alleles. For example, a matchup in which the recipient is inferior might be the following: the recipient genotype has three effectors each of which confer a 0.11 probability of killing a susceptible competitor genotype cell, while the competitor has three different effectors, each of which offer a 0.33 probability of killing a susceptible recipient genotype cell. A recipient genotype cell would therefore have a 33% chance of killing an adjacent competitor cell, while the competitor would have a 100% chance of killing an adjacent recipient cell.

Recipient genotype cells had a probability, ranging from 0-0.005, of replacing one of their T6SS effector protein / immunity pair alleles with one obtained from the environment each time step. This was performed in a weighted manner, according to the frequency of recipient vs. competitor alleles in the population, and did not depend on the spatial distribution of those cells in the population. Biologically, this is analogous to cells sampling DNA from a well-mixed bulk media above a surface-attached population. Note that this means that HGT can occur in both directions, with the recipient genotype replacing its T6SS with those from the competitor, and in future HGT events, reverting back to its original T6SS genotype. HGT events occur in a stepwise manner, so only one allele can be replaced per timepoint. The probability that adjacent cells succeed in killing was calculated as the sum of the per-allele killing probability for all of T6SS loci that differ between the cells. Simulations were run in Python (code available upon request). For Figure 4 and Supplementary Movies 2 and 3, fitnesses were calculated as the ratio of recipient:genotype cells at timepoint *t*, divided by their starting ratio. The fitness landscapes shown in these figures were calculated across a 20×20 factorial combination of HGT rates and competitor T6SS efficacies for 15,000 timepoints.

### Assessing fitness with variable competitors

We examined whether or not HGT could act as a hedge against unpredictable competitors by simulating pairs of interactions, in which the T6SS killing efficacy of the recipient strain varied symmetrically (*i.e*., ± 9.1, 27.2, 36.4 or 45.5% above and below) around the competitor’s killing efficacy. We calculated the geometric mean of fitness (raw fitness calculated as above) for these pairs of interactions across a range of HGT rates (0-0.005), with five replicates of the simulation run per treatment combination.

## Acknowledgements

This work was supported by the Gordon and Betty Moore Foundation grant #4308.07 and NSF grant #MCB-1149925 to B.K.H. and NSF grant #DEB-1456652, NASA Exobiology grant #NNX15AR33G, and a Packard Foundation Fellowship to W.C.R.

## Author contributions

JT, SSW, Conception and design, Acquisition of data, Analysis and interpretation of data, Drafting or revising the article; WCR, Conception and design, Acquisition of data, Analysis and interpretation of data, Drafting or revising the article, Funding acquisition; BH, Conception and design, Analysis and interpretation of data, Drafting or revising the article, Funding acquisition, Project administration.

